# Establishment of *Wolbachia* strain *w*AlbB in Malaysian populations of *Aedes aegypti* for dengue control

**DOI:** 10.1101/775965

**Authors:** W. A. Nazni, A. A. Hoffmann, A. Noor Afizah, Y. L. Cheong, M. V. Mancini, N. Golding, M. R. G. Kamarul, A. K. M. Arif, H. Thohir, H. S. Nur Syamimi, M. Z. Nur Zatil Aqmar, M. M. Nur Ruqqayah, A. Siti Nor Syazwani, A. Faiz, M. N. F. R. Irfan, S. Rubaaini, N. Nuradila, M. M. N. Nizam, M. S. Mohamad Irwan, N. M. Endersby-Harshman, V. L. White, T. H. Ant, C. Herd, H. A. Hasnor, R. Abu Bakar, M. D. Hapsah, K. Khadijah, D. Kamilan, S. C. Lee, M. Paid, K. Fadzilah, B. S. Gill, H. L. Lee, S. P. Sinkins

## Abstract

Dengue has enormous health impacts globally. A novel approach to decrease dengue incidence involves the introduction of *Wolbachia* endosymbionts that block dengue virus transmission into populations of the primary vector mosquito, *Aedes aegypti*. The *w*Mel *Wolbachia* strain has previously been trialed in open releases of *Ae. aegypti*; however the *w*AlbB strain has been shown to maintain higher density than *w*Mel at high larval rearing temperatures. Releases of *Ae. aegypti* mosquitoes carrying *w*AlbB were carried out in 6 diverse sites in greater Kuala Lumpur, Malaysia, with high endemic dengue transmission. The strain was successfully established and maintained at very high population frequency at some sites, or persisted with additional releases following fluctuations at other sites. Based on passive case monitoring, reduced human dengue incidence was observed in the release sites when compared to control sites. The *w*AlbB strain of *Wolbachia* provides a promising option as a tool for dengue control, particularly in very hot climates.

## Introduction

There are around 90 million symptomatic cases of dengue each year [1] with severe disease in around 1% of cases, including life-threatening haemorrhage or shock syndrome. Reducing abundance of the primary vector mosquito *Aedes aegypti* using insecticides and breeding site reduction remain the main strategies for dengue control, but are relatively ineffective. The introduction of *Wolbachia* endosymbionts into *Ae. aegypti* and demonstration of dengue transmission-blocking [2-6] has been followed by the use of the *w*Mel strain (from *Drosophila melanogaster*) for ‘population replacement’, resulting in this strain reaching and maintaining a high and stable frequency in north Queensland [7-9], where there has been a near-cessation of imported dengue outbreaks [10].

*Wolbachia* replacement involves the induction of cytoplasmic incompatibility (CI), a reproductive manipulation imposing a pattern of crossing sterility that provides an advantage to *Wolbachia*-carrying females. CI enables rapid spread to high frequency in insect populations once a threshold frequency has been exceeded, depending on host fitness parameters and CI / maternal transmission rates [7, 11, 12]. Different *Wolbachia* strains vary considerably in their effects on *Ae. aegypti* fitness parameters [13-16], and thus their population invasion / maintenance capacity (indeed strain *w*MelPop, a higher-replicating variant of *w*Mel with high fitness cost, was unable to maintain itself in *Ae. aegypti* populations [15]).

Recent reports suggested that *w*Mel may show reduced density and cytoplasmic incompatibility when *Ae. aegypti* larvae are reared at high temperatures [13-14, 16], which also matched temperatures previously recorded in wild *Ae. aegypti* larval sites elsewhere [17]; larvae in containers exposed to sunlight for part of the day would experience even higher temperatures than the recorded ambient air temperatures. However, *w*AlbB proved to be much less susceptible to the effects of similar high rearing temperatures [14, 16]. This suggests that *w*AlbB might be well suited for population replacement in hot environments, given its ability to effectively block transmission of dengue and other arboviruses [5]; *w*AlbB has not previously been deployed in this way.

The primary aim of these field trials was to assess whether *w*AlbB can be established / maintained at high frequency in urban *Ae. aegypti* in greater Kuala Lumpur, Malaysia, and what conditions are most conducive to establishment. In Malaysia over 100,000 dengue cases were reported in 2016, with an annual cost estimated at $175 million [18, 19]. Extended periods with daily peak temperatures exceeding 36°C occur in Kuala Lumpur. In light of *Wolbachia* strain difference in temperature responses, which may impact both the *Wolbachia* population frequency and the efficacy of dengue blocking in wild populations, this hot tropical environment provides an opportunity to test whether *w*AlbB-carrying *Ae. aegypti* with different effects on host fitness components to *w*Mel [14, 16] can invade an area where dengue is endemic. Information was also collected on dengue incidence in release areas and in matched control sites.

## Results

### Site selection

Intervention sites with persistent occurrence of dengue over the previous four years were selected (Table 1), in accordance with a WHO-recommended criterion for dengue intervention trial design [20]. Four primary intervention sites were chosen in Selangor State to represent different building types in order to explore *Wolbachia w*AlbB spread and maintenance dynamics in different settings (Table 1; Fig. S1): high-rise (18 floor) apartment buildings, 4/5-floor flats, 5 floor combined shop and flat buildings, landed terraces, and landed houses (Fig. 1). For two of the sites, releases were also conducted at adjacent smaller secondary sites with different building type. Where possible, boundaries to mosquito movement on at least a portion of the perimeter were incorporated in order to minimize immigration from surrounding areas - highways of six lanes and above / rivers / grassland and park areas [21]. Estimates of *Aedes* species composition and population size over time were obtained using ovitraps (Figs. 2, S2, S3). *Ae. albopictus* is present at all the sites (Fig. S3) (the latter is to be targeted subsequently for replacement releases using dengue-blocking strains [22, 23]. Community engagement is a very important component of *Wolbachia* transmission-blocking programs, given that biting female mosquitoes must be released into the environment; community consent and strong support for releases was obtained in all sites (see Materials and Methods).

**Table 1.** Information on release and control sites identified from criteria discussed in main text. *rate per 100,00, from Jan 2013 until start of intervention. Release sites are in bold.

| Type of site | Site name | Blocks (bl), floors (f) | Dengue Incidence* | Approx. no. households | Area (m<sup>2</sup>) |
| --- | --- | --- | --- | --- | --- |
| **Release-MH** | **Mentari Court** | **7bl, 18f** | **7,371.41** | **3,469** | **90,267** |
| Control-M1 | Lagoon Perdana | 7bl, 18f | 4,974.05 | 3,322 | 46,713 |
| Control-M2 | PJU 8 Apartment Flora Damansara | 8bl, 24f | 6,937.23 | 2,200 | 74,726 |
| Control-M3 | Seksyen 13 Apart. Brunsfield Riverview | 3bl, 14f | 11,725.74 | 532 | 10,329 |
| Control-M4 | Seksyen 13 Apartment Perdana | 4bl, 5-10f | 11,977.35 | 656 | 28,486 |
| Control-M5 | Desa Mentari | 11bl, 12-18f | 2,566.06 | 6,560 | 54,126 |
| Control-M6 | Kelana Puteri & Putera Condominium | 11bl, 16-20f | 7,726.39 | 1,322 | 56,586 |
| Control-M7 | Section U2 Apartment Ilham | 4 bl, 18f | 9,176.59 | 576 | 16,366 |
| Control-M8 | Section 7 Apartment Baiduri | 4 bl, 9-10f | 8,547.01 | 312 | 21,000 |
| **Release-SF** | **Section 7 Flats** | **51bl, 5f** | **11,539.09** | **2,930** | **276,261** |
| **Release-SL** | **Section 7 Landed** | **landed houses** | **14,592.93** | **248** | **113,817** |
| Control-S1 | Flat Section 6, 7 (BI52-58) & 8 | 42bl, 5f | 6,850.23 | 1,790 | 154,060 |
| Control-S2 | Flat Section 16, 17 & 18 | 68bl, 4-5f | 6,341.79 | 1,712 | 167,197 |
| Control-S3 | Flat Section 19, 20 | 37bl, 5f | 6,196.28 | 1,806 | 170,790 |
| Control-S4 | Flat Section 24 | 77bl, 5f | 3,504.36 | 4,260 | 151,620 |
| **Release-SC** | **Section 7 Commercial Centre** | **15bl, 4-5f shop apartment** | **13,325.22** | **1,408** | **119,025** |
| Control-C1 | Section 15 (Dataran Otomobil) | 18bl, 4-5f | 9,042.88 | 1,406 | 76,048 |
| Control-C2 | Flat Section 27 | 31bl, 4-8f | 4,715.82 | 3,100 | 157,364 |
| Control-C3 | Flat & Apartment Section U3 | 17bl, 5f | 3,677.25 | 1,800 | 104,763 |
| Control-C4 | Apartment Section U5 | 23bl, 5-7f | 3,681.08 | 2,736 | 139,404 |
| **Release-AL** | **AU2 Landed, Keramat** | **landed houses** | **4,170.03** | **1,239** | **727,548** |
| **Release-AF** | **AU2 PKNS Flats** | **18bl, 4f** | **1,452.66** | **672** | **50,598** |
| Control-A1 | AU5 Landed, Keramat | landed houses | 4,176.53 | 724 | 585,476 |
| Control-A2 | Taman Samudera & Taman Sunway, Batu Caves | landed houses | 4,270.45 | 1996 | 759,719 |
| Control-A3 | Bandar Tasik Puteri Fasa 6 & 7, Rawang | landed houses | 3,482.59 | 1675 | 841,977 |

**Fig. 1.**
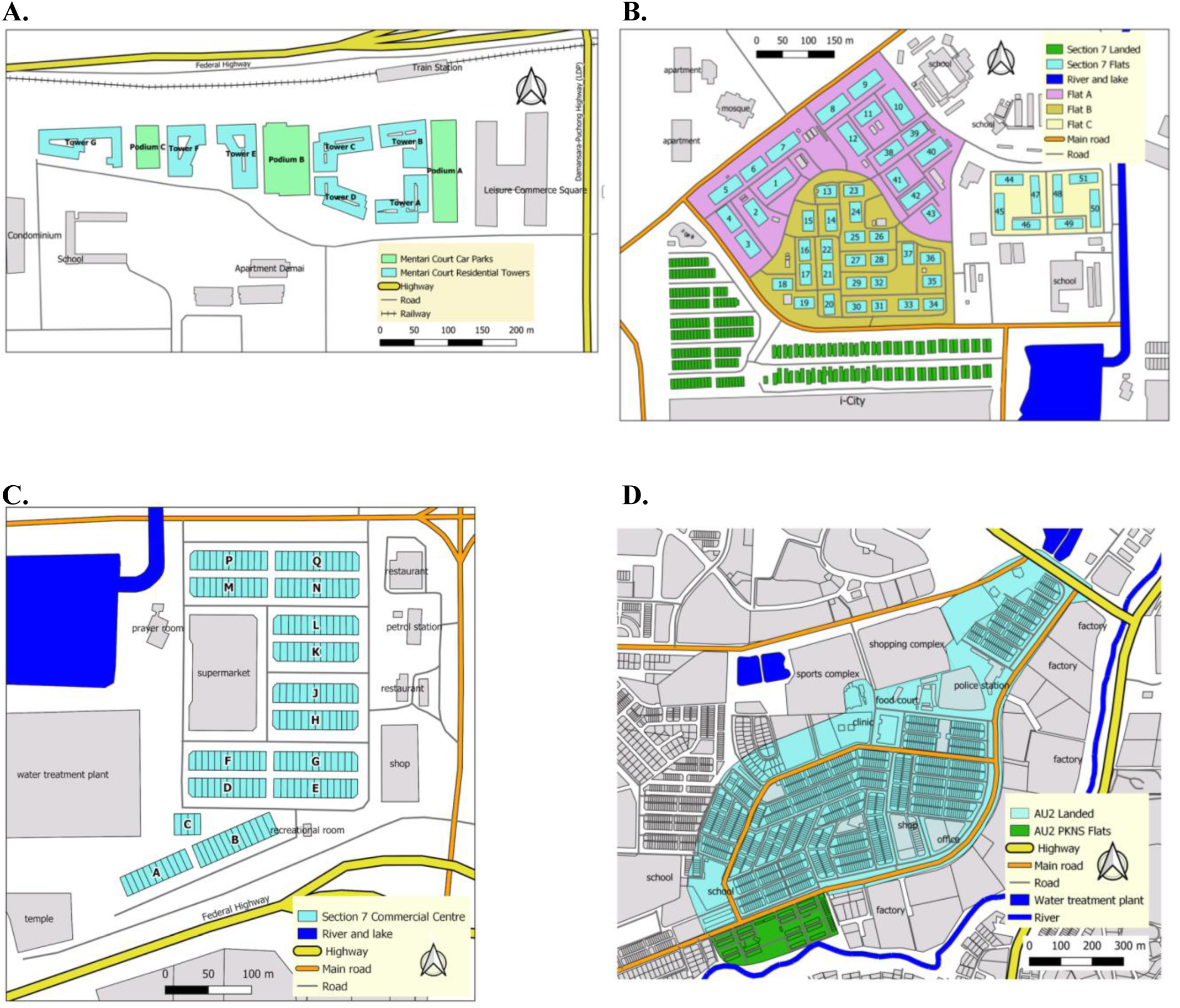
Maps of six release zones. **(A)** Mentari Court; **(B)** Section 7 Flats **and** Section 7 Landed; **(C)** Section 7 Commercial Centre; **(D)** AU2 landed **and** AU2 PKNS flats.

**Fig. 2.**
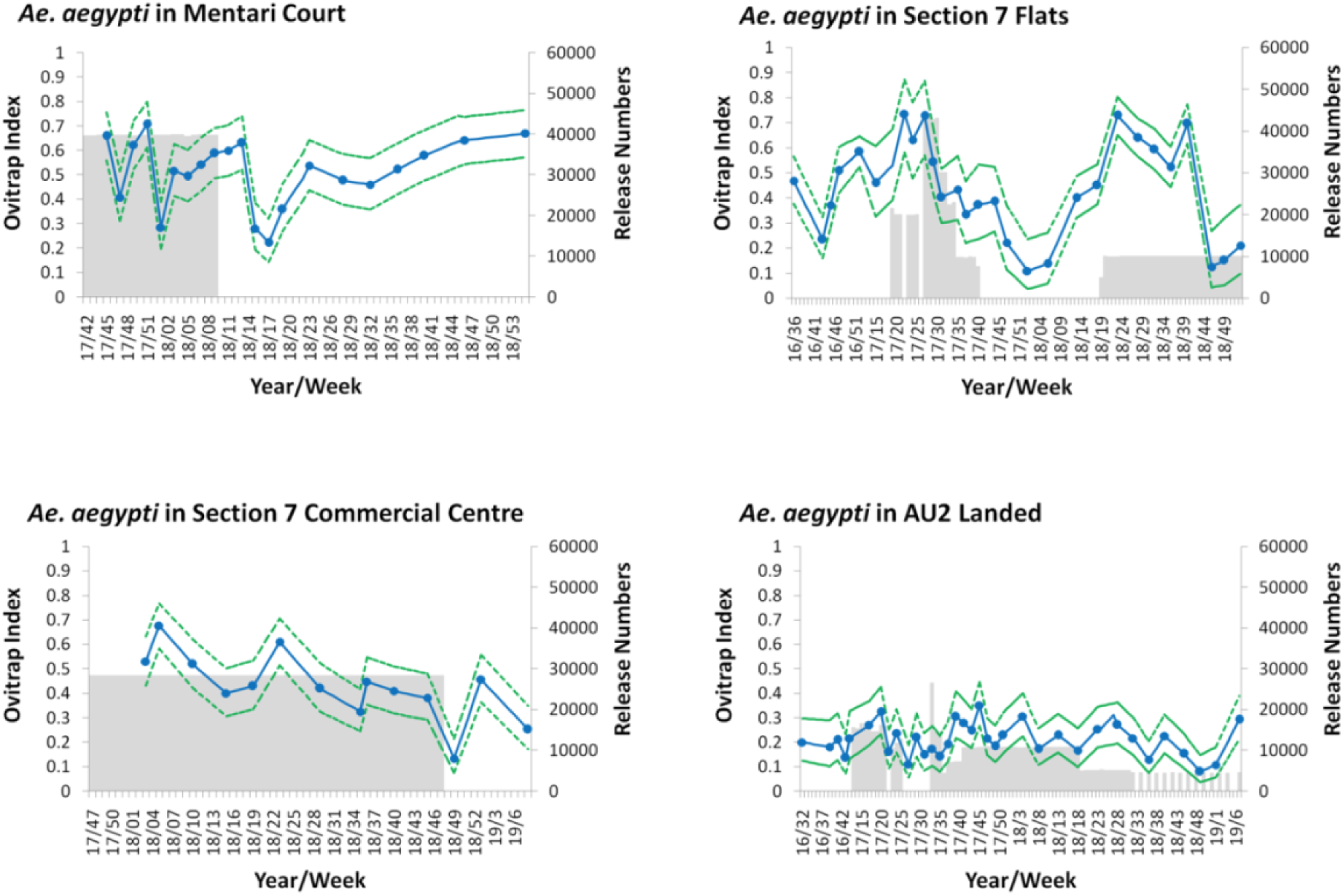
*Ae. aegypti* population size estimates at release sites. Ovitrap index (*Ae. aegypti*-positive traps divided by total number of traps) measured during the release/monitoring period. Grey shaded areas represent release periods; 95% confidence intervals are shown as dotted lines.

### Releases

In order to obtain a fit, locally adapted and competitive *Ae. aegypti w*AlbB line for release [16], four generations of backcrossing to field-collected local material were carried out. An important factor in relative fitness is insecticide resistance, given increasing levels of resistance [24] (particularly to pyrethroids). Susceptibility was compared between F1 adults from field-collected individuals and the release line using bioassays and found to be similar for pyrethroids, as well as to the organophosphates fenitrothion and pirimiphos (Fig. S4). The backcrossed *Ae. aegypti w*AlbB line was mass reared in preparation for releases. Wing measurements taken from mass-reared adults were in the range expected to produce fit, competitive release mosquitoes based on studies with Australian *Ae. aegypti* [25, 26] (average (SD) for males 2.28 (0.10) mm, females 2.96 (0.11) mm).

*Wolbachia* invasion depends on it exceeding a frequency dictated by fitness effects, incompatibility and maternal transmission; this was used to estimate required release rate with the ovitrap index (traps positive for *Ae. aegypti* divided by total traps per site) providing an estimate of population size [7]. Adult mosquitoes were released weekly in the morning on a pre-determined grid (with one cup of 50 mosquitoes released on a grid on ground / second floors in Section 7 and on every third floor in Mentari Court). After around 4 weeks of releases, *Wolbachia* frequency monitoring commenced using ovitraps, with the resulting eggs returned to the laboratory, raised to adults and a selection of the *Ae. aegypti* samples from each trap used for *Wolbachia* qPCR analysis [27]. In one of the sites, Section 7 Commercial Centre, eggs rather than adults were released: covered containers to which approximately 200 eggs in 400 mL water with larval food had been added (Fig. S5) were left out for 2 weeks for the adults to emerge on site.

### *Changes in* Wolbachia *and mosquito numbers*

*Wolbachia* frequency rose rapidly to over 80% at all sites (Fig. 3). Following the cessation of releases, the *Wolbachia* frequency has remained stable and high (98% 12 months after releases ceased) in Mentari Court. At the AU2 and Section 7 Flat sites, the frequency exceeded 95% but subsequently fluctuated following cessation of releases. Immigration of *Wolbachia*-free mosquitoes from surrounding untreated areas can reduce *Wolbachia* frequency where there is a low population size and relatively weak boundary barriers to mosquito population movement [21] which occurred at these locations; also in Section 7 Flats a 29-hectare construction site (‘i-City’), 159 m from the release zone, likely provided larval breeding sites and a source of wildtype *Wolbachia*-free migrants. Therefore, it was decided to resume releases at a lower release rate, whereupon *Wolbachia* frequencies rapidly rose again (Fig. 3).

**Fig. 3.**
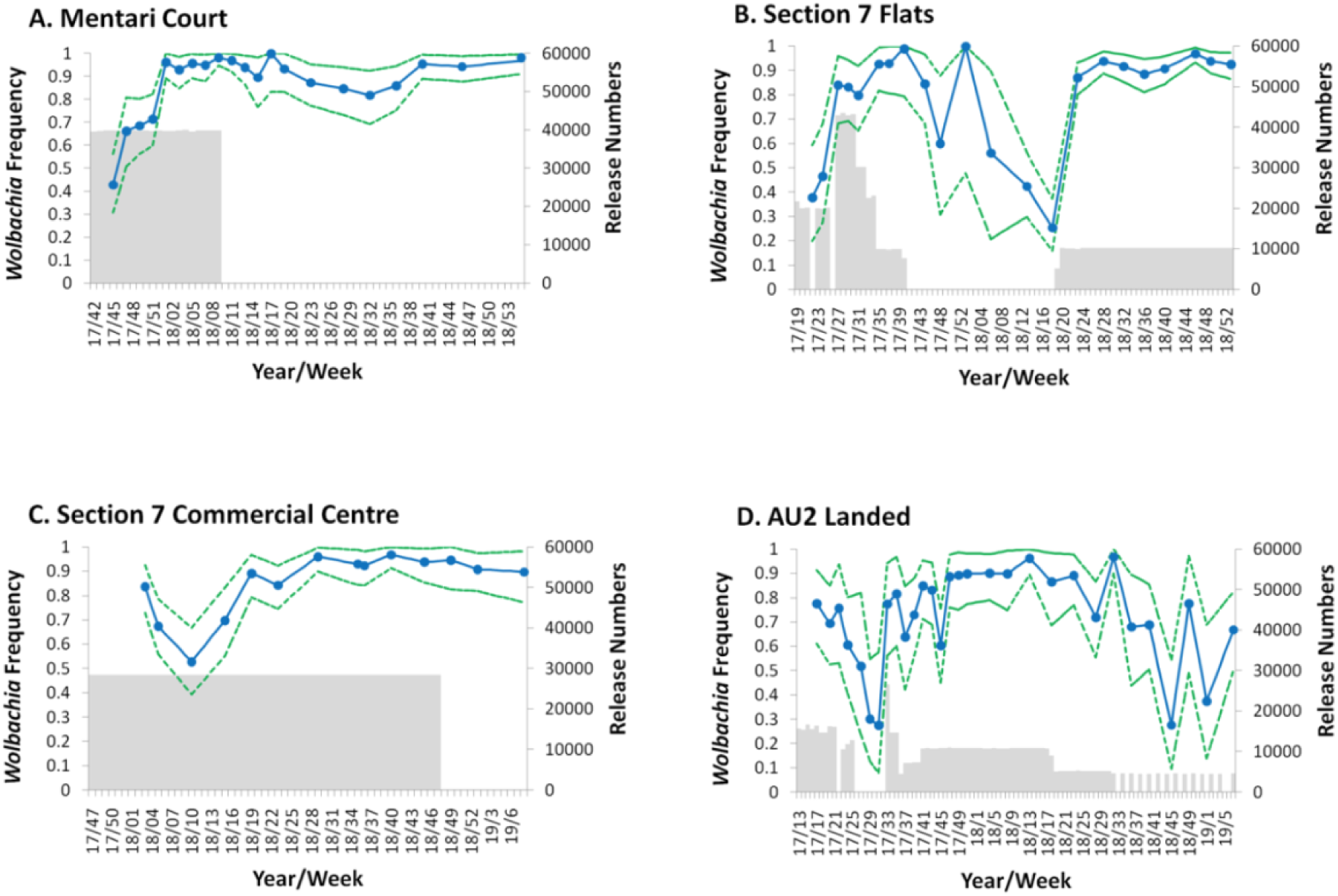
*Wolbachia* frequency during and after releases at six sites. Monitoring was conducted using ovitrapping and qPCR. **(A)** Mentari Court; **(B)** Shah Alam Section 7 Flats; **(C)** Shah Alam Section 7 Commercial Centre; **(D)** AU2 Landed. Release numbers are shown in grey shading; 95% confidence intervals are shown as dotted lines.

Population size monitoring before and during the releases (Fig. 2) also confirmed that there were no major population spikes associated with the release phase in the sites; this is expected given that mating between released males and wild females result in embryo death due to cytoplasmic incompatibility, balancing the population-increasing effects of female releases. In fact, in some sites post-*Wolbachia* establishment *Ae. aegypti* population density appeared to be lower than previously recorded based on ovitrap index (Fig. 2). More data are needed to support this observation, but the trend is consistent with a population suppressing effect caused by the fitness cost associated with *w*AlbB, particularly with respect to a steadily increasing mortality over time of desiccated eggs when added to water for hatching [16]. Areas with a higher proportion of temporary rather than permanent breeding sites, where periodic flushing of quiescent eggs is important, are predicted to experience a higher level of population suppression following invasion. Post-release community surveys revealed that a majority of residents did not notice any increase in mosquito biting (Fig. S6).

### Dengue incidence

Human dengue incidence from 2013-2019 were compared between release sites and matched control sites, based on data recorded by the Malaysian National Dengue Surveillance System. Control sites were selected for comparable dengue incidence to the release sites in the period from 2013 to the start of releases (Table 1); within the same District as the release site where possible, to ensure similar non-*Wolbachia* dengue control activities (except that a wider area was used for some of the Mentari Court sites, in order to ensure that some sites with similar or higher incidence compared to the release site could be identified); and building type of the ‘primary’ site in each location, to meet similar mosquito and human population characteristics (size, movement, etc.) (Table 1). Dengue incidence was reduced following releases in all intervention sites (Figs. 4, S7). A Bayesian time series model [28-30] produced an estimate of dengue case reduction of 40.3% over all intervention sites (95% credible interval 5.06 – 64.59) (Fig. S8), with posterior probability of a reduction in intervention sites post-release of 0.985.

**Fig. 4.**
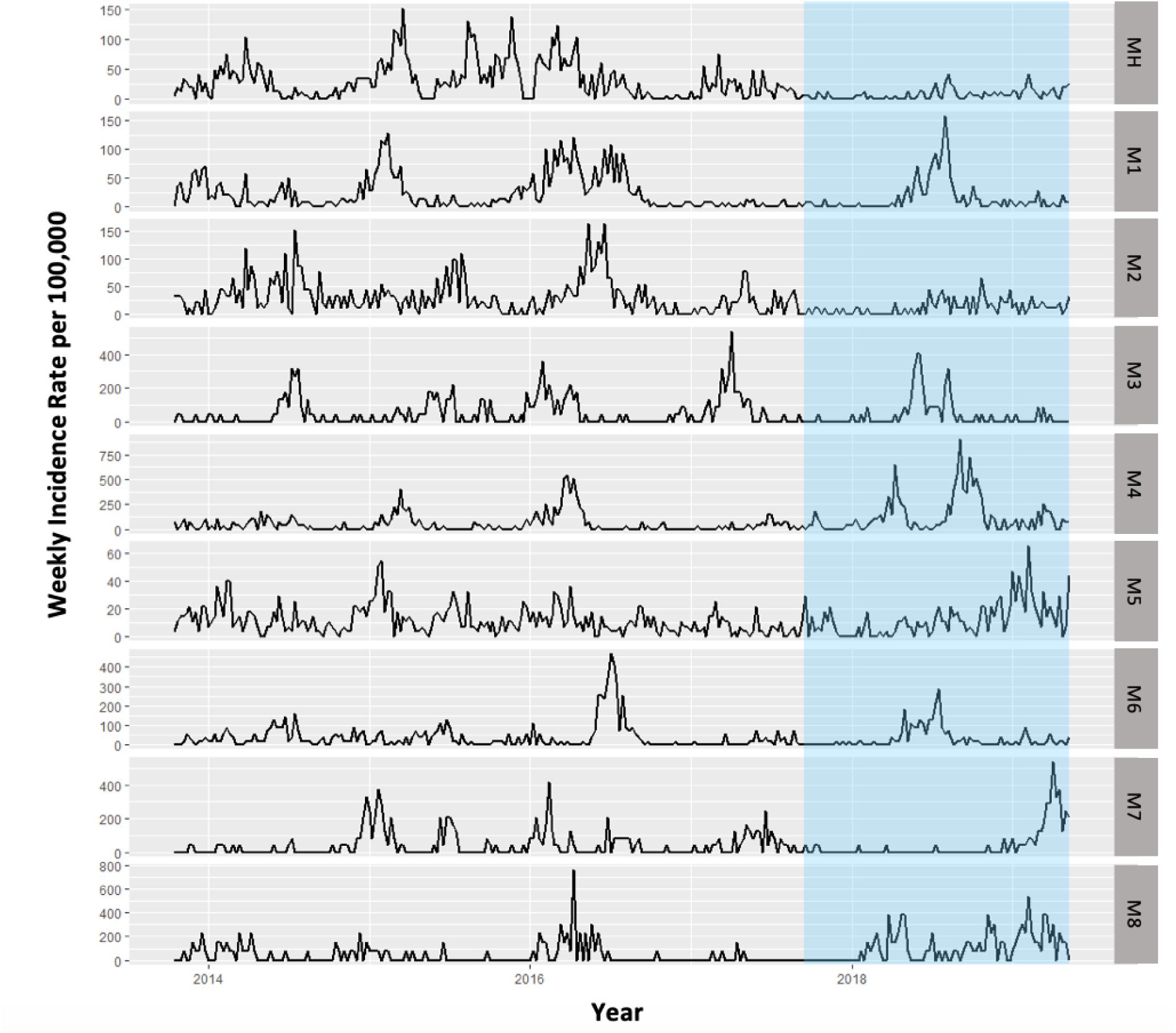
Dengue incidence from 2013 in Mentari Court (MH) and matched control sites (M 1-8). The period during and after commencement of *w*AlbB-carrying *Ae. aegypti* releases are indicated by blue (for other sites and their matched controls see Fig. S7). Incidence is calculated as total confirmed dengue cases per total population * 100,000.

In the 18-floor apartment buildings of Mentari Court, prior to *Wolbachia* releases dengue cases were high since at least 2013 (Fig. 4) despite intensive control efforts involving repeated rounds of clean-up ‘source reduction’ coupled with community engagement, and repeated thermal fogging (Fig. 5), none of which proved effective in reducing dengue incidence. The introduction of *Wolbachia w*AlbB has, in contrast, reduced dengue cases to a point where insecticide fogging by the local health authorities was no longer considered necessary (Figs. 4,5).

**Fig. 5.**
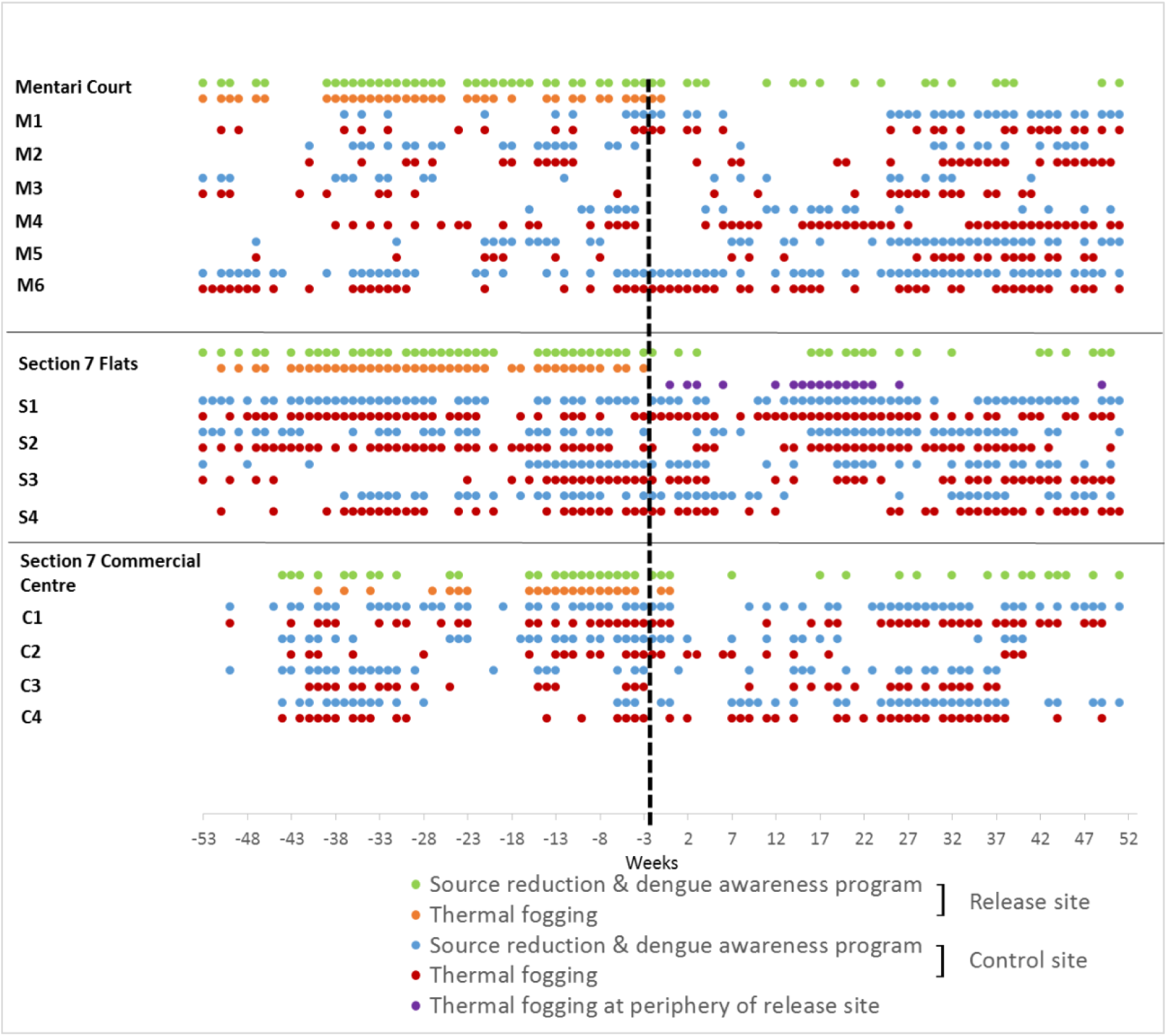
Dengue control activities (other than *Wolbachia* release) at the Section 7 Commercial Centre, Mentari Court and Section 7 PKNS Flats in release and matched control sites. The date of first *Wolbachia* releases is indicated by the black dashed line, and activities carried out in the year before and year after this date are shown. Data were only available for three release sites. Thermal fogging at the periphery refers to fogging done at nearby construction sites.

## Discussion

The results indicate the successful introduction of *w*AlbB *Wolbachia* into release sites in Selangor. The frequency of *w*AlbB has remained high at two sites following invasion (Mentari Court, Commercial Centre) and the frequency at Mentari Court is currently still >90%, two years after releases were terminated. An important consideration with respect to achieving stable high *Wolbachia* frequency is *Ae. aegypti* population size versus movement; in Mentari Court there is a relatively high population per unit ground area compared to the Section 7 zones, given *Ae. aegypti* were collected in ovitraps all the way up to the 18th floor. This will need to be incorporated into the design of wider strategy roll-out; for example buffer release zones should be incorporated around the perimeter of high-transmission localities that have small area, low *Ae. aegypti* population size or no clear natural boundaries to *Ae. aegypti* movement. The importance of area size in successful invasions has previously been recognized for *w*Mel introductions in Cairns, Australia, where *Wolbachia* failed to establish at one small site following releases despite successfully establishing at larger nearby release sites [21].

The successful deployment of egg containers for release at the Commercial Centre site has proven to be logistically much less laborious than adult releases and will also greatly facilitate wider roll-out. This reflects the fact that eggs produced in the laboratory can be easily cut into strips and transported to the field, where water and food can be added to containers. In contrast, adult releases require transfer of larvae to release containers where they can pupate and eclose, which requires additional handling of the immature stages by laboratory personnel. Egg containers were previously used in releases of *w*Mel in Townsville, Australia, where they also proved successful [10]. However, if density-dependent processes occur in such containers it could slow the rate of *Wolbachia* invasion [31].

The dengue incidence data points to a reduction in incidence at release sites following *Wolbachia* introductions when compared to control sites. Reduction to zero cases would not be an expected outcome even if the strategy is 100% efficient, given that *Ae. aegypti* bite during the daytime and thus dengue can be acquired outside of the place of residence (for example at work or school) during the day, and also given that *Ae. albopictus* is abundant at these sites (Fig. S3) and has yet to be targeted using *Wolbachia* replacement. Both factors make the effect of the intervention more difficult to detect with passive surveillance as was used here; only the effects of *Wolbachia* on local transmission of dengue by *Ae. aegypti* in the release zone are being detected in these comparisons. Nevertheless, clear differences in incidence between intervention and control sites were observed. The overall effect size produced by the Bayesian model of 40% dengue case reduction is thus a conservative estimate; active surveillance using seroconversion of naïve pre-school children would provide a more accurate measure of the effect of the releases on local transmission, given a much higher proportion of locally acquired cases are expected in this cohort [20], but this was beyond the budget and scope of the current study.

In summary, the results clearly demonstrate the capacity of *Wolbachia* strain *w*AlbB in urban *Ae. aegypti* to become established and maintain itself at high frequency in *Ae. aegypti* populations. Releases in a larger number of diverse intervention sites are now being undertaken in conjunction with the Malaysian Ministry of Health to comprehensively evaluate *w*AlbB effects on dengue transmission in different settings. The cessation of fogging in the release zones due to reduced dengue cases also points to the economic sustainability of the approach, given that large sums are spent annually on insecticides for dengue control [19]. Longer periods of dengue monitoring post-release will further increase the accuracy of the estimated effect size of the intervention on dengue incidence, and as more areas become *Wolbachia*-positive the proportion of imported cases will also fall. The establishment of *w*AlbB under high temperature conditions as reported here points to it being a promising option for deployment in very hot tropical climates. Transmission blocking for other viruses including Zika, chikungunya, and yellow fever could also be achieved using *Wolbachia* [4, 32, 33].

## Materials and Methods

### Backcrossing and mass rearing

To maximize the fitness and competitiveness of the mosquitoes to be released, backcrossing to freshly field-collected material was carried out. A total of 100 3-5 day-old females from the *w*AlbB *Wolbachia* line [16] were placed into 24 × 24 × 24 cm cages, together with the same number of 3-5 day-old males per cage of F1 / F2 mosquitoes from Shah Alam. After being left for mating for 2-3 days, they were allowed to blood feed on mice (Malaysian National Institute of Health approval number NMRR-16-297-28898). F1 females were then crossed to males of the field strain, again F1 / F2, to obtain the backcross one (B1) generation. The process was repeated twice to obtain generation B3. The presence of *Wolbachia* at 100% frequency was confirmed in the B3 generation by qPCR as described below.

Eggs were weighed on paper in 0.225 g lots (approx. 15000 eggs) and submerged in tap water (after 1 week of desiccation) in plastic containers (16 × 16 × 8 cm), after being exposed to an air vacuum to stimulate hatching. Eggs were then transferred to a 36 × 26 × 5.5 cm trays containing 1 L of tap water. Two days after hatching, L2 larvae from the 15000 eggs were sieved and transferred into 500 mL beakers filled with seasoned water (tap water stored overnight to dechlorinise). Aliquots of larvae were taken from the beaker using 10 mL plastic pipette tips (50 larvae per aliquoted mL). Ten aliquots (1 mL each) were placed into 36 × 26 × 5.5 cm trays filled with 1 L seasoned water. Cow liver powder (Difco, Becton, Dickinson and Co, Sparks, MD USA) and half-cooked cow liver were supplied to larvae daily at rates of .06, .08, .16, .31, .64, .32, .32, .32, .16, .08 and .06 mg per larva from days 1 to 11.

Pupae were transferred into small plastic containers with seasoned water and transferred into a mosquito cage (approximately 1000 pupae per cage). The mosquitoes were supplied with 10% sucrose solution with vitamin B complex. For blood feeding, two laboratory-reared mice were left in the cage overnight. Two days post-feeding, ovitraps (diameter 7 cm, height 9 cm) lined with A4 paper and with 200 mL of seasoned water were left for egg laying for four days. The paper was dried and then sealed in a plastic bag until use.

Wing measurements were carried out periodically on 15 mass reared males and 15 females for quality control. Both left and right wings of individual mosquitoes were dissected and measured under saline using DIMAS Professional version 5.0 (2) and an SSZ-T-3.5x mm graticule.

### Insecticide susceptibility assays

Adult bioassays for insecticide resistance were carried out using strains from field *Ae. aegypti* from Shah Alam and *Wolbachia*-carrying *Ae. aegypti*, together with a susceptible Kuala Lumpur laboratory strain that has been in culture for over 1000 generations. One commonly-used pyrethroid (permethrin) and two organophosphates (fenithrothion and pirimiphos methyl) were tested. For the field-derived samples, 20-100 larvae were sourced from ovitraps and reared for 1-5 generations in the lab. The *Wolbachia* mosquito colony was also derived from Shah Alam (see above). Larvae were raised on liver powder and 3-5 day-old adult females were tested.

The bioassays followed the WHO resistance test (WHO, 2016) using insecticide impregnated papers (Vector Control Research Unit, Universiti Sains Malaysia, Penang. 10-20 adult females were used for each of the three biological replicates. The adults were exposed to papers impregnated with permethrin 0.25%, pirimiphos methyl 0.25% and fenitrothion 1.0%. Two controls were also set up for each treatment. Knockdown was scored every 5 min during a 1 hr exposure period. After exposure, the mosquitoes were transferred into paper cups with cotton soaked in 10% sugar solution. Mortality was recorded after 24 hrs. Tests were repeated during the period that the colony was used for releases, except in the case of pirimiphos methyl that was tested twice.

### Community engagement

This strategy requires community engagement and consultation: acceptance and support of the community are essential for the releases. Printed educational materials containing clear information (e.g. leaflets, posters, buntings and banners) on *Wolbachia* were displayed and distributed. A website was established for the *Wolbachia* Malaysia Project linked at www.imr.gov.my/wolbachia. The target group to engage was the head of household and communities. To engage the public, interactions were initiated with local government / political / religious and community leaders, using information kits, workshops / roadshows, meetings / briefings, carnivals and home visits. A total of 40 stakeholders and community engagement activities were conducted in places of worship and community halls. Communities and Ahli Dewan Undangan Negeri (State Assemblyman) were also invited to the Institute for Medical Research to experience first-hand the science of *Wolbachia*. These activities were designed so that the communities would possess a sense of belonging to the programme, and feel that their involvement was recognized. Public meetings reinforced the ground-level support for the trials as reflected by a willingness of community members to participate in the program. This high level of support continued throughout the release periods.

Engagement by a team from IPTK (Institute for Health Behavioural Research) in all 6 release sites was conducted by undertaking lectures / talks, using brochures, undertaking advertised discussions with resident groups, posting banners & bunting, providing information on a website, as well as by sending messages to resident through WhatsApp, and SMS. There were between 4 and 17 activities at a release site including meetings, briefings, dialogue, carnivals and home visits (fewer activities were carried out at sites that were used later in releases as information about the releases spread). All activities involved local political leaders, community leaders and residents. The aim was to get the community’s trust in *Wolbachia* releases and the target of the community engagement team was at least 80% of the population having been exposed to the *Wolbachia* program through one or more of these approaches. The majority of the resident populations in three sites were tenants (65-70% Flat PKNS Section 7, 60-70% Mentari Court, and 80% in Section 7 Commercial Centre); tenants were in general less responsive and less involved in lecture activities and group discussion compared to homeowners. In the AU2 Landed, Section 7 Landed and AU2 Flats sites, none of the residents were tenants.

The IPTK team carrying out engagement continuously sought community feedback throughout the period and found that the 80% exposure target was not reached in some release sites. Hence the IPTK team, after discussing with the head of the blocks and apartments as well as Joint Management Board (JMB), decided at these release sites to obtain agreement for project implementation from community leaders and each community Joint Management Board.

In Shah Alam Section 7 Flats and Section 7 Landed sites, the vast majority of residents surveyed agreed to be involved in the *Wolbachia* mosquito release project: 99.6% (650 of 652) of responding households gave their approval for the project. In AU2 Landed and AU2 Flats, 98.4% (501 of 509) of responding households gave their approval to the project. In Shah Alam, of those responding to the survey 62.9% were male and 27.5% female, while in AU2 Keramat, 50.7% were male and 33.2% female; those responding by WhatsApp and URL were unidentified and accounted for 9.7% and 16.1% from Shah Alam and AU2, respectively. In Mentari Court, the target group involved 20 community leaders rather than residents because attendance at resident meetings was low. All community leaders agreed to the release of *Wolbachia* in all the blocks and car parks in Mentari Court. The community engagement in Section 7 Commercial Centre involved briefing community leaders on the *Wolbachia* project by the IPTK team. After engagement, all 14 community leaders agreed for Commercial Centre Section 7 to be included in the project. The community leaders requested *Wolbachia* videos and health promotion materials for distribution to businesses, shops, houses and other premises, and this was provided.

For all release sites, the *Wolbachia* mosquito release progress was shared with the communities by inviting community representatives to the Institute on several occasions. An engagement activity in the community was also run where the development of *Wolbachia* carrying *Aedes aegypti* larvae were monitored in egg release breeding containers directly exposed to sunlight, partially shaded areas and shaded areas.

Feedback surveys on *Wolbachia* interventions were conducted prior to and after releases. Questions included the public perceptions to *Wolbachia* releases and dengue, the perception of the mosquito population size before, during and after the releases, the level of confidence in *Wolbachia* reducing dengue cases, opinions on breeding sites reduction after *Wolbachia* releases and level of concern about dengue transmission. Approximately a year after release initiation, the communities from the release sites were again invited four times for briefings and updates on the progress of the programme.

### Mosquito releases

A risk assessment was completed and permission was obtained for *Wolbachia* releases after being examined by NIH and unconditionally approved by the Malaysian Medical Research and Ethics Committee (MREC) (reference number KKM/NIHSEC/P16-566).

The areas of the 6 release sites are provided in Table 1 and maps of the sites are provided in Fig. 1. The dates for releases in Section 7 Flats, Section 7 Landed, Mentari Court, and Commercial Centre were 13 May 2017, 13 May 2017, 16 October 2017 and 20 November 2017 respectively, while the dates for AU2 Landed and AU2 Flats were 28 March 2017 and 13 September 2017 respectively. The 1st monitoring was conducted after 4 weeks of adult releases. However, for the egg releases in the Commercial Centre, the first monitoring was conducted after 4 egg releases, after 8 weeks on 15 January 2018.

Two days prior to the initial releases of *Wolbachia*-carrying *Ae. aegypti* in each site, fogging to suppress the wild populations was carried out using ultra low volume spray (ULV) of pyrethroids or organophosphates, in accordance with the Standard Operative Procedures of the Ministry of Health. During the trial release period, no fogging or space spraying activities were conducted within the release sites. GIS maps of release sites were prepared in order to construct release point grids, using the GIS software ArcGIS v10.5 and QGIS 3.2.3 Bonn. Release site maps were sketched based on the topography and cadastral map provided by the Department of Survey and Mapping Malaysia and with reference to Google Maps and 1MalaysiaMap (http://1malaysiamap.mygeoportal.gov.my/).

Mosquitoes were transported to the field sites in a van kept at ambient temperature. At Mentari Court, a total of 40,800 *Wolbachia*-carrying adult mosquitoes were released weekly in blocks A to G including indoor parking areas. The mosquitoes were released on every third level in each block at 4 release points, except for block G which had 6 release points. For the indoor parking building, there were 4 release points on the ground floor and 2nd floor. Four paper cups (diameter 8 cm, height 11 cm), each with 50 mosquitoes (mixed sex) were released. Following each release 10 containers were brought back to the laboratory and survival of adults was monitored; very little mortality (never > 5%) was observed. Releases were conducted for 20 weeks and terminated when the *Wolbachia* frequency reached >90% on 3 consecutive monitoring periods. At the final monitoring period 98% were *Wolbachia*-carrying, as assessed through ovitraps. At Section 7 Flats, 20,400 *Wolbachia*-carrying adults were released weekly in blocks 1 to 51. There were 8 release points across 2 levels (ground floor and 2nd floor), with 1 cup per release point. Releases were initially conducted across 26 weeks when *Wolbachia* frequencies exceeded >90% on 3 consecutive monitoring periods. Releases ceased for 4 months, and then a second release period was undertaken for 31 weeks at a lower rate (10,200).

At the Section 7 Commercial Centre, releases involved eggs rather than adults. Egg containers (14.5 cm x 12.5 cm) contained 150 *Wolbachia*-carrying eggs in 400 mL of water with 180 mg liver powder. A single release hole (2 cm) was covered with a stopper. Egg containers were placed at a release point in the shade, on mailboxes or near staircases. Containers were left for 2 weeks. After 5 days, the stopper was removed to allow mosquitoes to emerge. Hatching rate was assessed on 5 containers by placing them in a sealed container left in the field house. Adults were counted (75-80% of the eggs produced adults).

### Monitoring mosquito population size and Wolbachia *frequency*

Ovitraps were used to assess *Wolbachia* frequencies and to monitor numbers of *Ae. aegypti* and *Ae. albopictus*. Each ovitrap consisted of a plastic container (96 mm height, 67 mm diameter) with 150 mL water and a wooden paddle (2 cm x 7 cm). In Mentari Court, 100 ovitraps were set up on the ground. In the apartment buildings, these were placed on the ground floor, the 2nd, 5th, 8th, 11th, 14th and 17th floors. For the car park building, the 1st floor and 3rd floors were monitored. Ovitraps were collected after a week and the paddle+water was transferred to a plastic container (12 × 12 cm). All emerging mosquitoes were identified to species and a maximum of 10 *Ae. aegypti* per trap were used for *Wolbachia* screening.

In Section 7 Shah Alam, monitoring was done after 4 weeks from the first release using 183 ovitraps, with 3 ovitraps set up per block on the ground, 3rd and 5th floors. Initially, monitoring in Mentari Court and Section 7 Shah Alam was conducted every two weeks and later every month. In Section 7 Commercial Centre, 100 ovitraps were set on the ground, middle and top floors as evenly as possible across the release zone. Monitoring was initiated after the 4th egg release on a monthly basis.

Adults from ovitraps were stored in absolute ethanol at −80°C. DNA was extracted from individual mosquitoes using the Chelex® 100 resin (Bio-Rad Laboratories) method. Mosquitoes were homogenised in 175 µL of 5% Chelex® solution using TissueLyser II machine (Qiagen) and with 5 µL of proteinase K (20 mg/mL) (Bioron Life Science). The extraction was incubated in thermocycler at 65°C for 1 hour, followed by incubation for 10 minutes at 90°C_16_. *Wolbachia* was detected by high-resolution melting polymerase chain reaction (qPCR-HRM) [27] with 1:10 diluted DNA using the following *wAlbB1*-specifc primers: *wAlbB1*-F (5’-CCTTACCTCCTGCACAACAA) and *wAlbB1*-R (5’ – GGATTGTCCAGTGGCCTTA), as well as universal mosquito primers: *mRpS6*_F (5’-AGTTGAACGTATCGTTTCCCGCTAC) and *mRpS6_*R (5’-GAAGTGACGCAGCTTGTGGTCGTCC), which target the conserved region of the *RpS6* gene, and *Ae. aegypti* primers *aRpS6-*F (5’-ATCAAGAAGCGCCGTGTCG) and *aRpS6-*R (5’-CAGGTGCAGGATCTTCATGTATTCG), which target the *Ae. aegypti-*specific polymorphisms within *RpS6* and do not amplify *Ae. albopictus*.

Reactions were run as 384-well plates in a LightCycler 480 II (Roche). qPCR-HRM was performed in 10 µL reactions containing 2 µL of DNA, 0.08 µL of 50 µM forward+reverse primer, 2.92 µL Milli-Q water and 5 µL Ronald’s Real-Time Buffer (3.28 µL Milli-Q water, 0.4 µL MgCl_2_ (50 mM), 1.0 µL ThermoPol Reaction Buffer with 20 mM Magnesium (10x), 0.25 µL HRM Master (Roche), 0.064 µL dNTPs (25 mM) and 0.01 µL Immolase ™ (20 U/µL). qPCR was run following cycling conditions: 95°C for 10 min, followed by 50 cycles of 95°C for 10 seconds, 58°C for 15 seconds, 72 °C for 15 seconds. High resolution melting was performed by heating the PCR product to 95°C, and then cooling to 40°C. Then the temperature was increased to 65°C. Samples were considered positive for *Wolbachia* when the Tm for the amplicon produced by the *Ae. aegypti* primers was at least 84°C and the Tm for the *Wolbachia*-primer amplicon was around 80°C.

### Dengue control activities

In Mentari Court, many different control activities and education programmes have been carried out to suppress dengue. Fig. 5 shows that various activities such as source reduction, thermal fogging and Ultra Low Volume space spraying were conducted. Despite this activity, dengue cases remained high in this site. After the introduction of *Wolbachia*, there was a decrease in other activities undertaken to suppress dengue following a decrease in cases. Therefore, community engagement around *Wolbachia* decreased rather than increased other control activities at this site. At the other release sites, there was no increase in non-*Wolbachia* dengue suppression activities following the initiation of releases.

### Dengue incidence

Matched control sites were selected based on similarity of constituent buildings (at least 7 blocks of 12-20 floors for Mentari Court controls, at least 35 blocks of 4-7 floors for Shah Alam Flat controls, at least 15 blocks of 5-7 floors for Commercial Centre controls); and comparable dengue incidence between 2013 and the start of intervention periods. Notified dengue cases in release and control sites were recorded daily from local clinics and hospitals. All notified cases were confirmed using NS1 and IgM/IgG Combo Rapid Test Kits (RVR Diagnostics Sdn. Bhd., Subang Jaya, Malaysia) according to manufacturer’s protocols, based on the established Malaysian Dengue Clinical Practice Guidelines (CPG), available on http://www.moh.gov.my.

All notified dengue cases are recorded into the National e-dengue database at the District Health Office. Dengue cases in the population residing at the study sites were detected via this National Dengue Surveillance System. All cases were diagnosed using the National case definition guidelines (Case Definitions for Infectious Diseases in Malaysia 2017). A confirmed case of dengue was defined as fulfilling the clinical criteria for dengue infection with the following laboratory confirmation: detection of Dengue Non-Structural Protein 1 (NS1) from serum; and, detection of dengue IgM and /or IgG from a single serum sample. While there are limitations to all laboratory diagnostic tests for dengue, the dengue rapid test kits (RTK) applied in this National Surveillance System represent the most cost-effective point of care diagnostic testing at a population level. The standardized diagnostic criteria applied through this system ensures no biases in test results between the intervention and control sites.

### Bayesian time series model

A Bayesian time series model was used to estimate the reduction in dengue cases resulting from releases. The model structure was as follows:

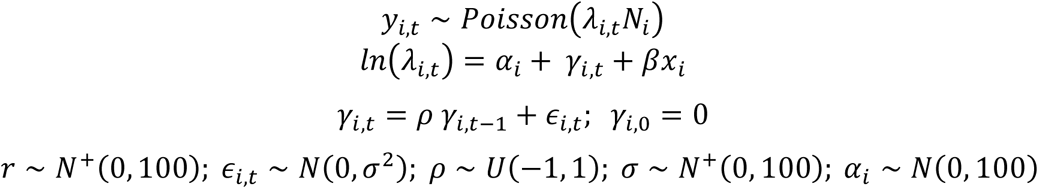

where the number of cases *y* at each site iand week *t* were assumed to follow a Poisson distribution, with the expected count given by the product of population at that site, and the per-capita incidence which varied varying through time and between sites. Each site had a separate time series of log-incidences *α*_*i*_+ *γ*_*i,t*_ with temporal correlation driven by an autoregressive model of order one, with parameters *ρ* and *σ*^2^ shared by all sites. Each observation therefore had a separate (temporally-correlated) random effect on the log scale, to account for extra-Poisson dispersion and temporal correlation. The intervention effect was represented by a parameter *β* and an indicator variable *x*_*i*_ for whether the observation was post-release at a release site. Model parameters were assigned vague normal; positive-truncated normal; or uniform priors. The model was used both to estimate the impact of the releases, and to assess the evidence from the dengue case data that releases lead to a reduction in incidence - quantified as the posterior probability that *β* is negative, in the absence of prior knowledge about the direction of the effect.

Posterior samples of model parameters were simulated by Hamiltonian Monte Carlo in greta [29] with 4 chains, each yielding 4000 posterior samples of model parameters after a warmup period of 1000 iterations during which period the leapfrog step size and diagonal mass matrix parameters were tuned. The number of leapfrog steps was sampled uniformly from between 30 and 40 throughout. Convergence was assessed by the Gelman-Rubin 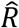 diagnostic, using the coda R package [30] (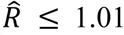 for all parameters) and visual assessment of trace plots. Model fit was assessed by posterior predictive simulation: a random dataset of *y*_*i,t*_ values was generated according to each posterior sample of *p*_*i,t*_ and *r*, and the distributions of the simulated *y*_*i,t*_.values were compared with the observed *y*_*i,t*_. The analysis code is freely available online. (code attached with submission and archive URL to be provided here on acceptance).

## Acknowledgments

The primary funding for the study was provided by Wellcome Trust Award 108508, with additional funding from Wellcome Trust Award 202888 and National Health and Medical Research Council of Australia Awards 1118640 and 1132412.

## Author contributions

S.P.S., A.A.H. conceived the study; W.A.N., A.N.A., Y.L.C., M.V.M., M.R.G.K., A.K.M.A., H.T., H.S.N.S., M.Z.N.Z.A., M.M.N.R., A.S.N.S., A.F., M.N.F.R.I., S.R., N.N., M.M.N.N., M.S.M., N.M.E-H., V.L.W., T.H.A., C.H., H.A.H., R.A.B., M.D.H., K.K., D.K., S.C.L., M.P., K.F. performed the experimental work; A.A.H., Y.L.C., N.G., B.S.G., M.V.M., S.P.S. analysed the data; A.A.H., S.P.S., W.A.N., H.L.L. supervised the project; S.P.S., A.A.H. wrote the manuscript; all authors reviewed the manuscript.

The authors declare no competing interests

## Supporting Information

**Fig. S1.**
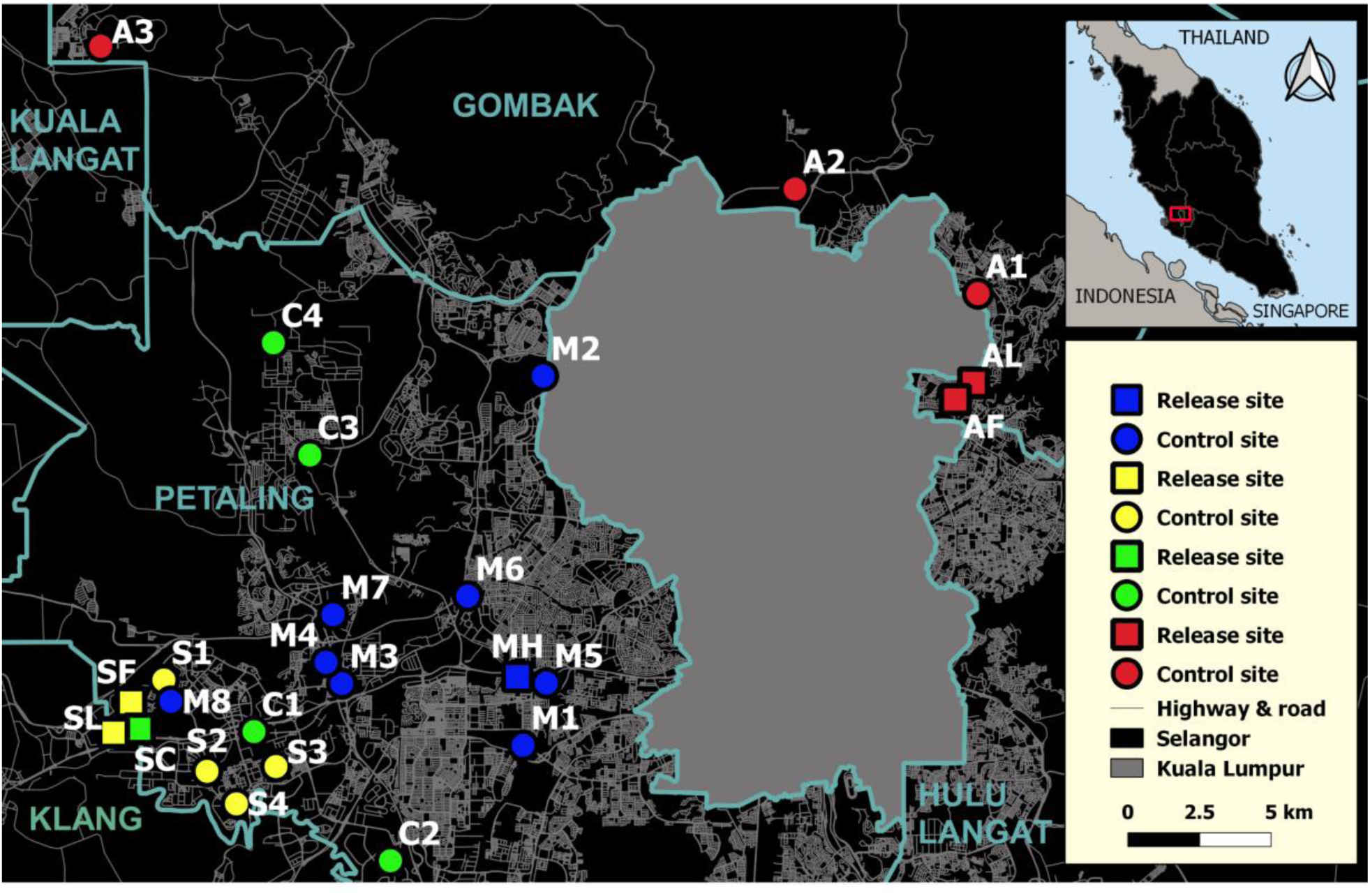
Maps of all six release zones and controls in the Petaling and Gombak Districts of Selangor: Mentari Court high rise (MH) and controls (M1-M8); S7 Flats & Landed (SF & SL) and controls (S1-S4); S7 Commercial (SC) and controls (C1-C4); AU2 landed & AU2 PKNS Flats (AL & AF) and controls (A1-3). Table 1 contains additional details about release and controls sites. Map constructed using MapCruzin.com, OpenStreetMap.org under the Open Database License (http://opendatacommons.org/licenses/odbl/1.0/)).

**Fig. S2.**
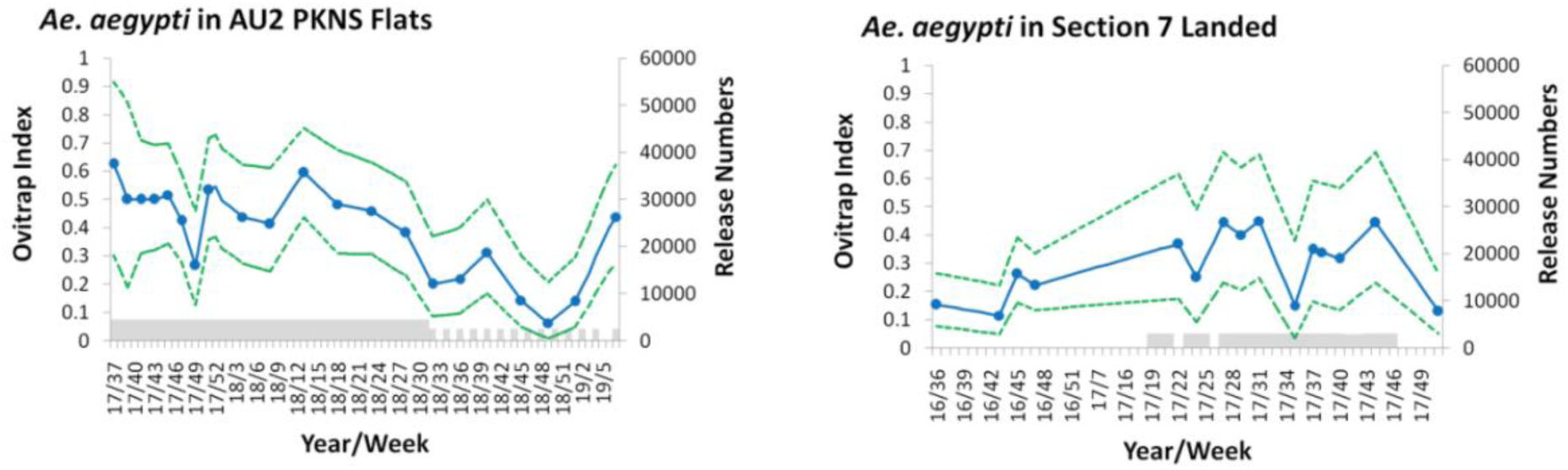
*Aedes aegypti* population size estimates for two smaller secondary release sites through ovitrap index (*Ae. aegypti*-positive traps divided by total number of traps) during the release/monitoring period. Grey shaded areas represent release periods; 95% confidence intervals are shown as dotted lines.

**Fig. S3.**
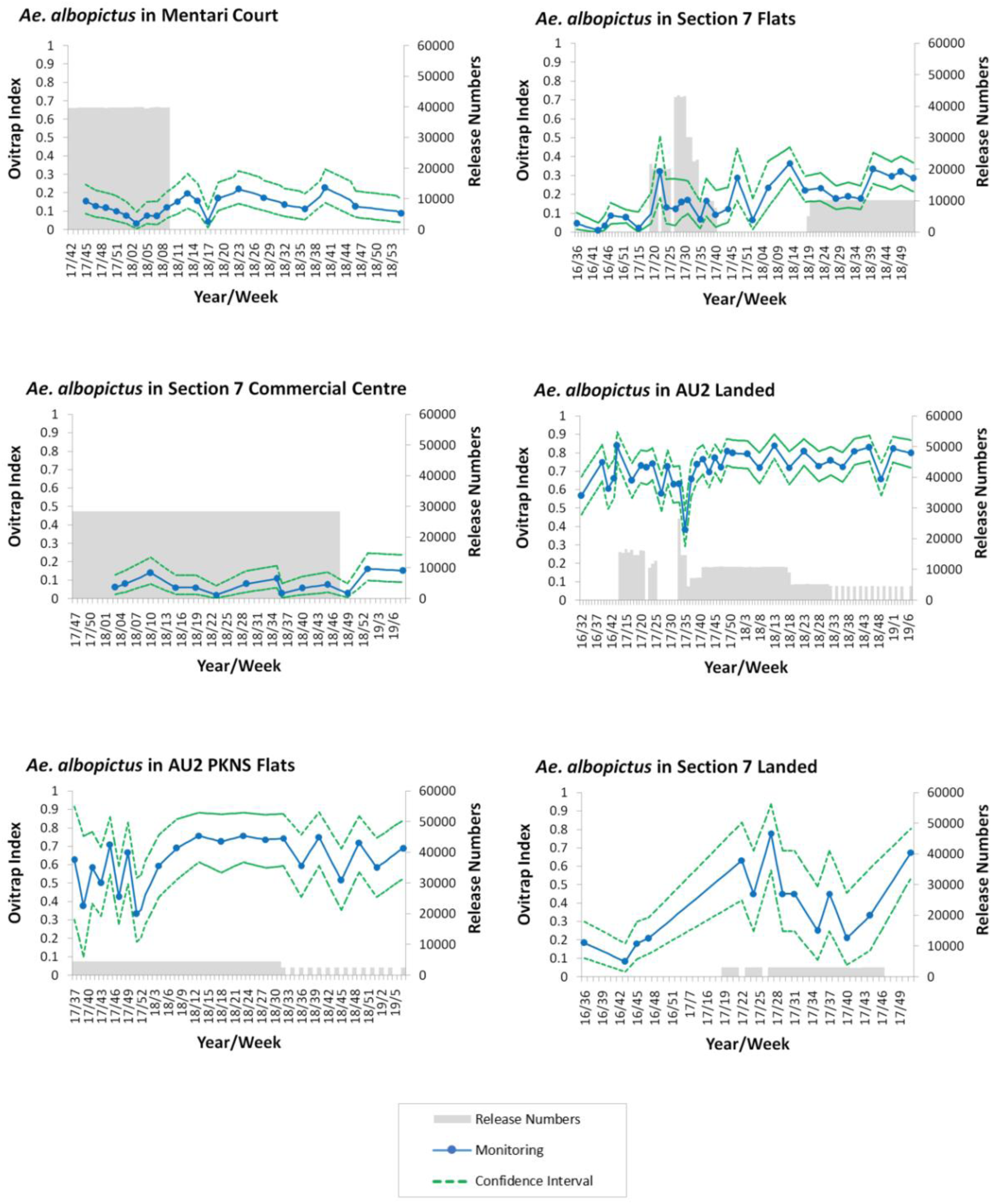
*Aedes albopictus* population size estimates at release sites measured by ovitrap index (*Ae. albopictus*-positive traps divided by total number of traps) during the release/monitoring period. Grey shaded areas represent release periods; 95% confidence intervals are shown as dotted lines.

**Fig. S4.**
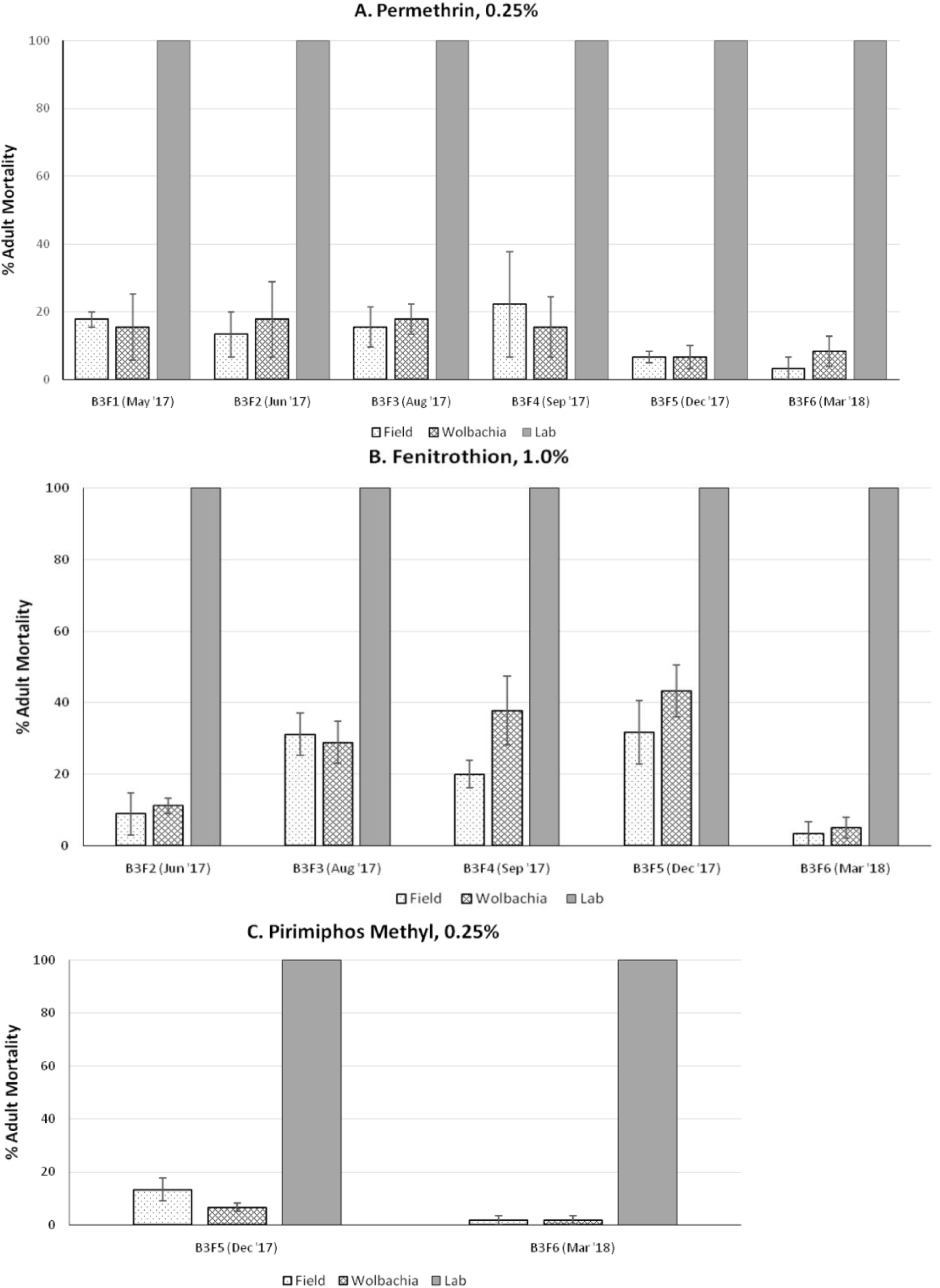
Insecticide susceptibility of the release line (“Wolbachia”) compared with the field-derived wild-type population of *Ae. aegypti* (“Field”) and a laboratory insecticide susceptible line (“Lab”) tested against three pesticides: permethrin, fenitrothion, and pirimiphos methyl. Bioassays measured adult mortality at discriminating doses. Error bars represent standard errors.

**Fig. S5.**
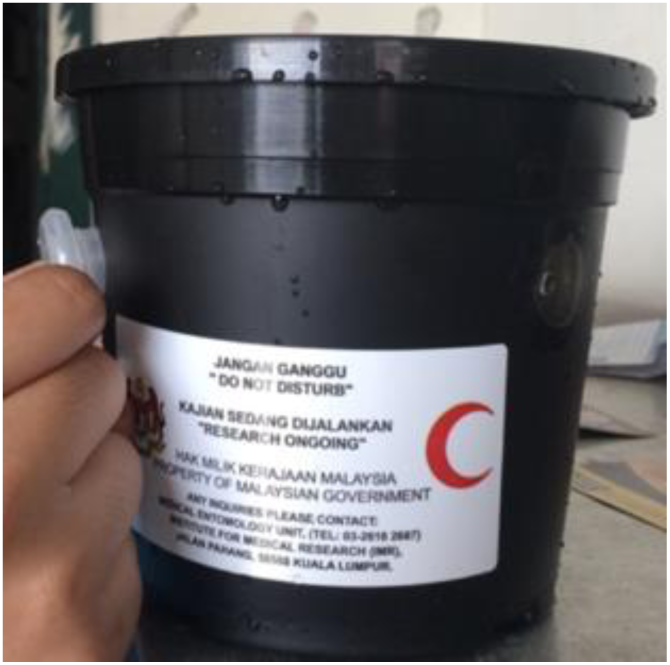
Example of an egg release container as used in the Section 7 Commercial Centre site; note the 2 cm diameter release holes and plug, removed to allow escape of adult mosquitoes.

**Fig. S6.**
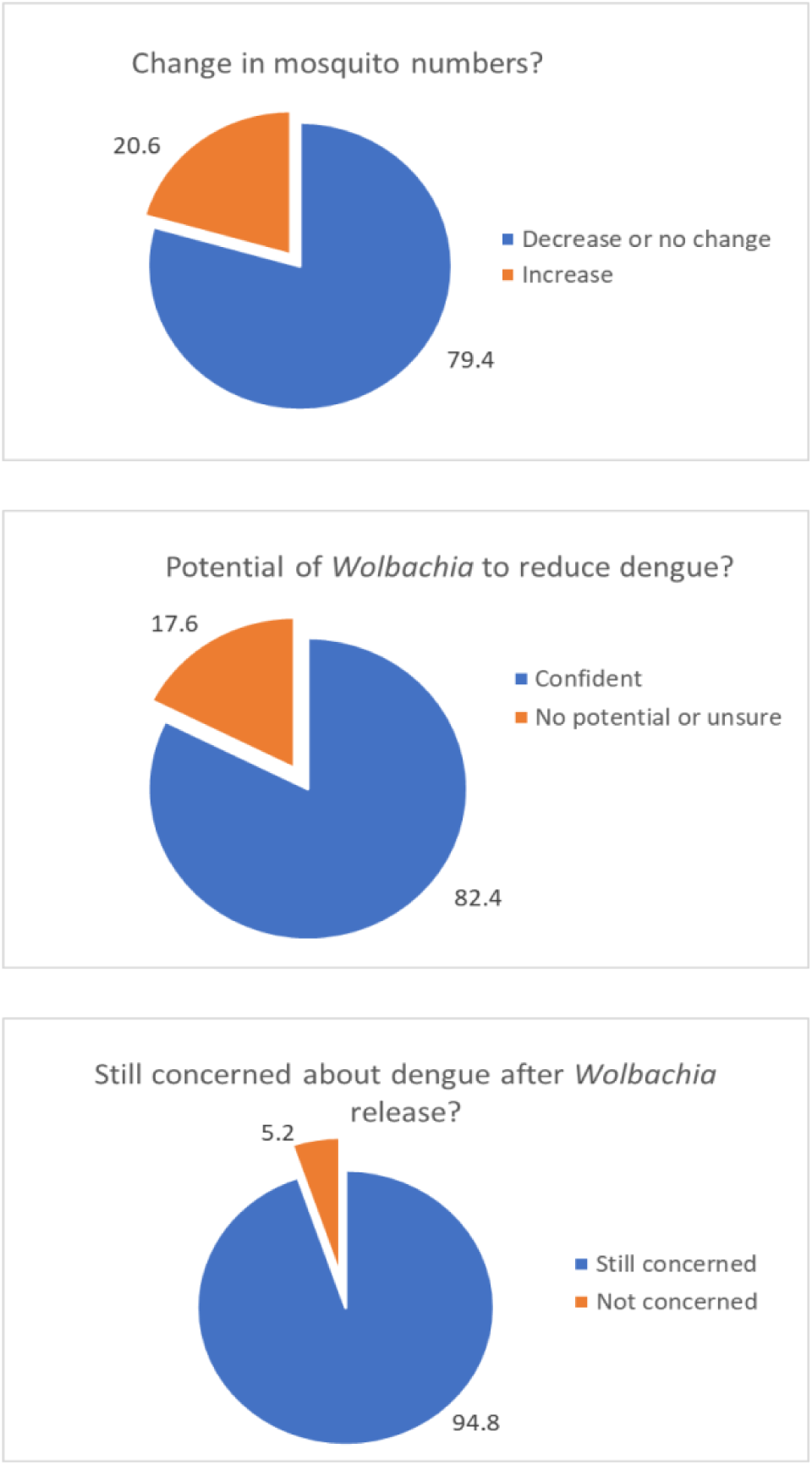
Post-release survey questionnaire results from community engagement activities in Section 7 Flats. Data were collected from a door-to-door survey in the Section 7 Flats release area 2 months after the beginning of the releases. 196 respondents were included, consisting of 40% females, with age classes 29-39 (15%), 40-50 (65%) and >60 (10%).

**Fig. S7.**
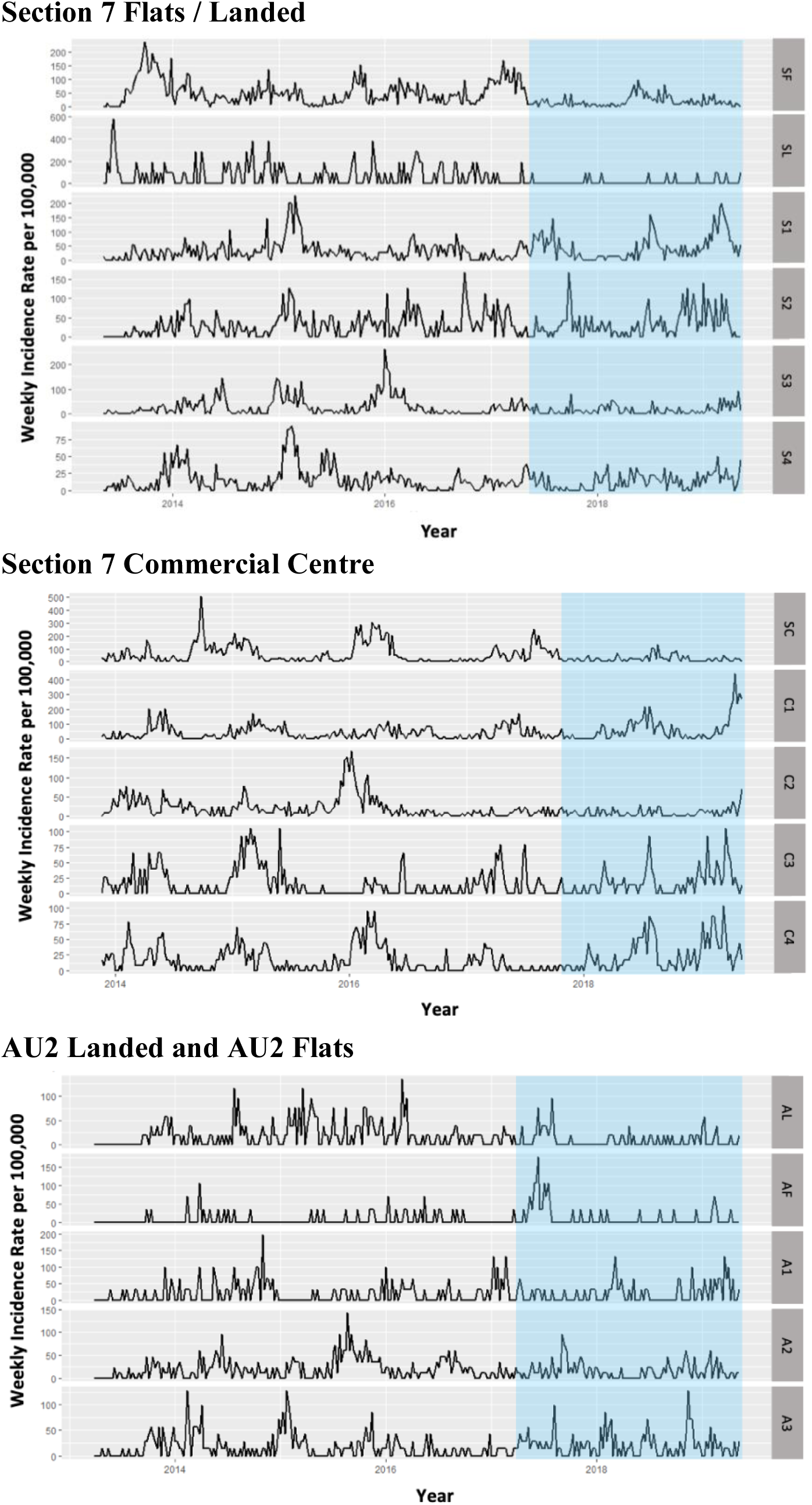
Dengue incidence plots at the Shah Alam Section 7 PKNS flats & Landed, Section 7 Commercial Centre, and AU2 Landed / Flats release and matched control sites. SF, SL, SC, AL and AF represents release areas on different sites. See Table 1 for additional details on control sites. The periods during and after commencement of *Wolbachia* releases are indicated by blue regions.

**Fig. S8.**
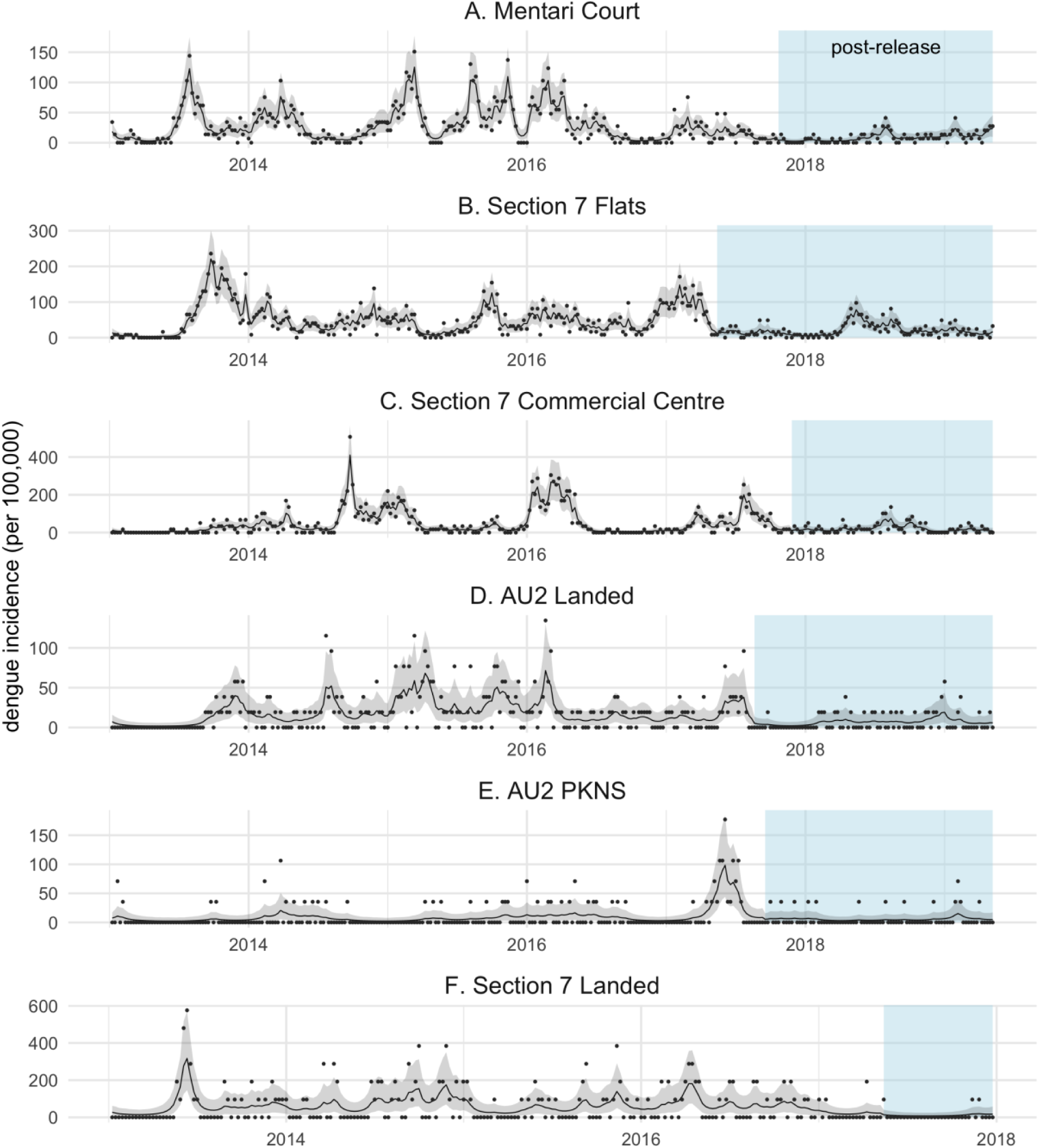
Dengue reduction following *Wolbachia* releases. Incidence of confirmed human dengue cases for each week of the study period in the release sites. Black lines and grey shaded areas show the posterior mean and 95% credible intervals of the incidence inferred from a Bayesian time series model. Points represent empirical incidences calculated directly from case data. The period during and after commencement of *Wolbachia-*carrying *Ae. aegypti* releases are indicated by blue regions.

